# Common Genetic Variants of The Cardiac Sodium Channel Alter Patient Response to Class 1b Antiarrhythmics

**DOI:** 10.64898/2026.01.14.699482

**Authors:** Martina Marras, Joshua D. Josef, Thomas Schuster, Heather L. Struckman, Emily Wagner, Lucas Lee, Kyle J. Smith, Janice M. Amsler, Lucy S. Woodbury, Erica Marquez, Jonathan D. Moreno, Jennifer NA Silva, Jonathan R. Silva

## Abstract

Several common variants have been identified in SCN5A, which encodes the cardiac sodium channel α-subunit Nav1.5 and is targeted by class I antiarrhythmics. Lidocaine and its analog mexiletine both have a primary amine that blocks Na^+^ current. While lidocaine is highly effective in terminating ventricular tachycardia after acute myocardial infarction, mexiletine has been shown to prevent arrhythmia induction in only ∼20% of patients. The factors underlying this inconsistent drug response are unclear. Here, we use cardiomyocytes that are derived from induced pluripotent stem cells to observe that a common polymorphism in the SCN5A gene, S1103Y, exhibits an altered pharmacological response to mexiletine, with enhanced use-dependent and tonic block of peak sodium current. In addition, an unexpected increase in late sodium current causes action potential prolongation. This paradoxical proarrhythmic phenotype shifts the paradigm of conventional antiarrhythmic therapy with mexiletine, suggesting that background variants may alter pharmacological responses leading to unanticipated consequences. Our results suggest that the unique genetic background of patients should inform therapeutic approaches to treat and prevent arrhythmias associated with common cardiac pathologies.

## Introduction

Cardiac rhythm disorders affect between 1.5 to 5% of the general population, and may result in malignant ventricular arrhythmias and sudden cardiac death^1^. Although the substrates and phenotypes of arrhythmias are varied, the vast majority are treated with antiarrhythmic drug therapies which target cardiac ion channels^2^. The voltage-gated sodium channel Nav1.5, encoded by the SCN5A gene and responsible for the rapid upstroke of the action potential and impulse propagation, remains a prominent target for antiarrhythmic drug therapy and molecules that target it comprise the Vaugh-Williams class I antiarrhythmic drug family^3^. Sodium channels activate rapidly upon membrane depolarization, followed by fast inactivation during which the channel is non-conductive, although still open, resulting in a narrow Nav current window^4–6^. Genetic or acquired anomalies that alter this tightly regulated gating are associated with cardiac conduction diseases and arrhythmias^7,8^. Despite decades of research into the basic mechanisms of Na^+^ channel blockade via local anesthetic drug action, multiple failed clinical trials (notably, the Cardiac Arrhythmia Suppression Trial (CAST) in 1991)^9^ and clinical experience confirm our inability to fully predict in which patients these drugs may be of benefit, and emphasizes the urgent need to systematically understand patient-specific characteristics that may alter drug efficacy^10^. To that end, personalized pharmacogenetic medicine is already being implemented in cardiology clinical practice, in the context of genetically focused antiplatelet testing^11,12^.

In addition to tonic block, which occurs when the drug reaches and binds to the receptor while the pore is still closed, several sodium channel blockers also exhibit use-dependent or frequency-dependent inhibition, indicating that the drug preferentially reduces channel conductance when it is in the open or inactivated state^13^. Specifically, class 1b antiarrhythmics, including lidocaine and mexiletine, are considered weak sodium channel blockers and favor binding to the open and inactivated states^14,15^. Lidocaine is a prototypical class 1b antiarrhythmic molecule and is prescribed to suppress ventricular arrhythmias either as monotherapy or often as adjunctive therapy with amiodarone^16^. Because of high first pass hepatic metabolism, lidocaine requires intravenous administration and thus is not used outside the acute, inpatient setting^16,17^. Consequently, mexiletine was developed as an oral analog of lidocaine^3,4,18^ and received FDA approval in 1969. Although mexiletine is orally available and has a relatively low toxicity after extended use, patient tolerability due to dose dependent gastrointestinal side effects also reduces clinical usage^14^.

Like lidocaine, mexiletine has been shown to preferentially inhibit persistent sodium current (31.5% reduction) compared to peak sodium current (10.5% reduction)^19^, which is beneficial to treat heart rhythm pathologies associated with gain-of-function SCNA mutations, such as long QT syndrome (LQTS) type 3. However, significant variability in mexiletine effectiveness has limited widespread adoption, with several reports of inconsistent results among individuals who carry different LQTS-associated SCN5A variants^19,20^. In-vitro studies have confirmed a similar pattern: despite a clinical phenotype of ventricular arrhythmia connected to increased late Na^+^ current, efficacy of mexiletine appears to be mutation specific. While LQTS-linked mutations are rare, several more common single-nucleotide perturbations have been identified in the SCN5A sequence, providing background genetic variability without inducing pathogenicity^21^. We hypothesize that common Nav1.5 polymorphisms, which appear with relatively high prevalence in the general population, may play an outsized role in determining mexiletine efficacy.

At present, the association between common variants and mexiletine response remains elusive. Some single-nucleotide SCN5A polymorphisms are present with more frequency in certain ethnicities. For example, S524Y and S1103Y appear in 6% and 13% of African-descendent populations^21,22^, respectively, and R34C is found in 9% of Blacks and 3% of Asians^14^. Other variants, like H558R, appear with high prevalence (∼9-29%) in the general population across all ethnicities^21,23^. In this study, we investigate whether common sodium channel polymorphisms that are not directly pathogenic modulate the mexiletine response, probing the link between genetic variation and drug effectiveness. The results demonstrates that the presence of S1103Y homozygosity produces an unexpected drug phenotype, with potentially proarrhythmic consequences. This study provides further insight into the differences in antiarrhythmic drug responses based on common SCN5A variants that are likely to be underappreciated, highlighting the importance of understanding the complete genetic makeup of patients to increase the success of clinical outcomes.

## Methods

### Plasmids

The human SCN5A [Genbank accession No. AC1377587] and SCN1B (Genbank accession No. A0A6Q8PHF7) genes, encoding the cardiac sodium channel α- and β-subunits respectively, were subcloned into a mammalian expression vector, pIRES, to allow high efficiency co-expression of both proteins from the same bicistronic mRNA transcript^24^. Site-directed mutagenesis for the R34C, S524Y, H558R and S1103Y variants was achieved using overlap extension polymerase chain reaction, followed by Gibson assembly and electrochemical transformation to introduce the mutated plasmid into Mega-X bacterial ultracompetent cells. Cells were cultured overnight in Luria Broth until they reached an optical density value of 1.5-2, measured as the ratio of absorbance at 260nm and 280nm with a NanoDrop spectrophotometer (ThermoFisher). NucleoSpin Plasmid Miniprep kits (Macherey-Nagel) based on silica membrane technology were used to extract and purify DNA. All mutations were confirmed with next-generation Sanger sequencing (Azenta).

### HEK Cell Culture and Transfection

Human embryonic kidney (HEK) 293 cells were used to assess the Nav1.5 electrophysiological properties of variant channels. 35mm cell culture dishes were seeded with ∼150,000 HEK293 cells to a confluency of ∼50-60% and maintained in Dulbecco’s Modified Eagle’s Medium supplemented with 10% Fetal Bovine Serum and 100 µM penicillin-streptomycin, in 37°C, 5% CO_2_ incubator. 24 hours after seeding, cells were transfected with one of the four β-pIRES-Nav1.5 plasmids using JetOptimus transfection buffer and reagent (Polyplus). The reaction included 600 ng of DNA, 60 µL of buffer and 0.6 µL of reagent per 35 mm-dish. Transfected HEK293 cells were incubated at 37°C for 36 to 48 hours. An EVOS cell imaging system was used to monitor the health and cell confluence daily. Before recording, cells were rinsed twice with Hank’s Balanced Salt Solution, no calcium, no magnesium, no phenol red (Gibco) and then singularized with a mild enzyme, Accumax (Live Cell Technologies), for 5 minutes at 37°C. After quenching the enzyme with growth medium, cells were allowed to recover at 4°C for one hour. Cells were resuspended with equal volumes of growth medium and standard electrophysiology external solution (135 mM sodium) to a concentration of ∼500,000 cells/mL in preparation for automated patch clamp recordings.

### hiPSC Culture and Differentiation

Human induced pluripotent stem cell derived cardiomyocytes (hiPSCs) from a wild-type donor (WTc-11 line) were obtained from the Human Cells, Tissues and Organoids Core at Washington University in St. Louis (WashU) and cultured as previously described^25,26^. Briefly, they were seeded on 6-well tissue culture plates pre-coated with 10% Matrigel (Corning) at a density of ∼30,000 cells per well and maintained in mTeSR+ medium (StemCell Technologies) at 37°C, 5% CO_2_. Fresh culture media was exchanged every 2 days according to manufacturer’s recommendations. Cells were passaged when they reached ∼90% confluence using ReLeSR dissociation reagent (StemCell Technologies), mTeSR+ and 1:1000 Rock inhibitor Y27632 (Tocris, 5μM). Differentiation into cardiomyocytes was achieved using small-molecule Wnt inhibitors according to published protocols. Briefly, iPSCs at ∼80% confluence were switched to RPMI medium supplemented with B27-insulin (B27i), containing 1:1000 CHIR99201 (Tocris, 6μM) and 1:1000 L-ascorbic acid (150 μg/mL, Sigma) (day 0). On day 2, the medium was replaced with RPMI/B27i containing 5 μM Inhibitor of Wnt Production 2 (IWP2) and 1:1000 (150 μg/mL) ascorbic acid. On day 4, the medium was replaced with RPMI/B27i containing 1:1000 (150 μg/mL) ascorbic acid. Spontaneously beating cardiomyocytes were observed starting at day 6 and kept in culture in RPMI with B27 complete (B27C) until day 20. Cells were metabolically purified with two rounds of lactate, subsequently singularized with trypsin and replated as thin monolayers, maintained in culture in RPMI/B27C^27,28^. All experiments were performed between day 30 and day 40.

### hiPSC Reprogramming and Genome Editing

Peripheral blood mononuclear cells (PBMC) were collected from a patient carrying S1103Y heterozygosity upon written consent. Frozen PBMC were expanded and reprogrammed into human induced pluripotent stem cells (hiPSCs) at the Genome Engineering & Stem Cell Center (GESC) at WashU and named MVET-55. Karyotyping and pluripotency staining was conducted for all clones. All work with hiPSCs was approved by the institutional review board #MVET IRB 202007184 at WashU. S1103Y^+/+^ iPSCs were generated from both WTc and MVET-55 iPSCs at the GESC. S1103Y^-/-^iPSCs were also generated from the MVET-55 iPSCs as control. Briefly, a single guide RNA (sgRNA) sequence (5’-actggcctcggcctcagagGagg-3’) was designed to target exon 18 of SCN5A and substitute serine with tyrosine (tcc > tAc) at locus 1103. The sgRNA and Cas9 enzyme were complexed into a ribonucleoprotein and delivered to cells via nucleofection. Non-homologous end joining was used to knock in the S1103Y mutation. Positive clones were analyzed via next generation sequencing, sorted, and underwent final quality control assays (short tandem repeat profiling, mycoplasma testing, genotyping and karyotyping).

### Electrophysiology

The Patchliner (Nanion Technologies), a fully automated planar patch system, was used for electrophysiological recordings. The external standard solution contained 140mM NaCl, 4mM KCl, 1mM MgCl_2_, 2mM CaCl_2_, 5mM D-glucose monohydrate, and 10mM Hepes, titrated with NaOH to pH 7.4, osmolarity 298mOsm. The internal solution contained 10mM CsCl, 10mM NaCl, 110mM CsF, 10mM EGTA, 10mM Hepes, titrated with CsOH to pH 7.2, osmolarity 285mOsm. Inward sodium currents were elicited with 250-ms depolarizing pulses from a holding potential of −100mV to +50mV, in 10mV increments. The steady state inactivation (SSI) was measured with 250ms pulses from −150 mV to +20mV, in10mV increments, followed by a test pulse to −20 mV and an inactivating step to −100mV. Conductance-voltage (GV) and SSI curves were quantified by fitting data to a Boltzmann function (Eq. 1):

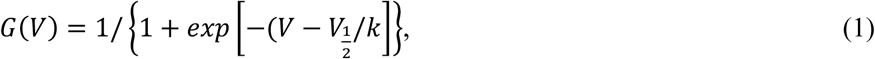

where V_1/2_ is the half-activation voltage and k is the slope factor.

The channel recovery from inactivation was measured with depolarizing pulses from −120mV to +20mV at varying time increments ranging from 2 to 1000 msec. The data was fit with a double exponential function to quantify both fast and slow phases of recovery (Eq. 2):

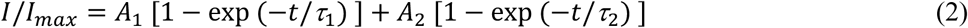

where I is the peak sodium current, I_max_ is the maximal peak current, A_1_ and A_2_ are the proportional coefficients, t is time, ρ_1_ and ρ_2_ are the fast and slow recovery time constants, respectively. The late component of the sodium current was measured at 20ms from the peak current measurement and reported as the ratio between late and peak values.

To characterize the pharmacological response to mexiletine of wild type and variant channels, cells were held at –100 mV and then depolarized with a conditioning pulse to −20 mV to induce sodium channel opening. For use dependent block, 50 consecutive pulses were applied at 10Hz frequency. The first pulse of each sweep was considered an approximation for 0 Hz, or tonic block. Serial dilutions of mexiletine in standard external solution (1 µM, 5 µM, 10 µM, 100 µM, 500 µM, 1 mM and 2 mM) were applied acutely during each sweep. The dose response curve was obtained by fitting the data to a log(inhibitor) vs normalized response equation with variable slope in Prism/Graphpad. From the dose response, the half maximal inhibitory concentration (IC_50_) was calculated to assess the potency of each drug in wild type and variant sodium channels. For each cell, the relative inhibition of current amplitude was normalized against the peak current value recorded after the first depolarizing pulse to account for variability in cell size and current density. 95% confidence intervals, unpaired t-tests for comparisons within each group, and one-way ANOVA for between-groups comparisons were used to determine statistical significance, with error bars representing SEM and p < 0.05 considered significant (Prism/GraphPad).

### Fluorescence microscopy

Action potentials were recorded optically. Cell monolayers were externally labeled with a fluorescent voltage sensitive dye (BeRST-1, 0.75 μM, Lumencor) 15 minutes before recording and paced at 1Hz (MyoPacer, IonOptix) under an optical microscope (Nikon ECLIPSE Ts2R-FL) to capture APD. As the voltage propagates across the cell membrane, fluorescence intensity changes are used as reporters of the AP wave. Signals were analyzed on Matlab using an automated open-source waveform analysis software developed by the Huebsch lab at Washington University in St. Louis^29^, and normalized for fluorescence intensity. Statistical significance between each group was assessed using an unpaired t-test, with error bars representing SEM and p < 0.05 considered significant.

### Fluorescent Immunolabeling

Singularized hiPSC-CMs with MVET-55 background (3 lines: S1103Y^-/-^, S1103Y^+/-^, S1103Y^+/+^) were plated at 40,000 cells/plate on 35mm glass bottom dishes (#1.5). Samples were fixed with 2% PFA (5 minutes at room temperature [RT]), washed with PBS (3 x 10 minutes at RT), and permeabilized 0.2% Triton X-100 in PBS (15 minutes at RT). hiPSC-CMs were then incubated with blocking agent (1% bovine serum albumin, 0.1% triton in PBS for 2 hours at RT). For subdiffraction confocal imaging, samples were labeled with primary antibodies (overnight at 4°C) and washed in PBS (3 × 5 minutes at RT). After primaries, samples were labeled with secondary antibodies (2 hours at RT) and washed in PBS (3 × 5 minutes at RT). Proteins of interest were labeled with well-validated custom and commercial antibodies: anti-sarcomeric alpha actinin antibody (mouse monoclonal antibody, Cat. #MA1-22863, Invitrogen), N-cadherin (N-cad) (mouse monoclonal antibody, Cat. #33-3900, Invitrogen), and cardiac voltage-gated sodium channel 1.5 (NaV1.5) (a previously validated custom rabbit polyclonal antibody)^30^. Samples labeled for confocal microscopy were labeled with goat anti-mouse and goat anti-rabbit secondary antibodies conjugated to Alexa 488 and Alexa 568 (1:4,000; ThermoFisher Scientific).

### Confocal Imaging

Confocal images of singularized hiPSC-CMs (Figure 2) were acquired using a Leica Sp8 Lightning single photon confocal microscope equipped with 5 solid-state lasers (405, 488, 514, 552, and 638 nm, 12 mW each), a 63x/1.4 numerical aperture oil immersion objective, 2 HyD GaAsP detectors, and 2 high-sensitivity photomultiplier tube detectors (Leica). Images were collected as single planes snapshots and with Nyquist sampling (or greater) as previously described^31^.

## Results

Local anesthetics, including class 1b antiarrhythmics lidocaine and mexiletine, inhibit sodium channels via two distinct mechanisms, either by permeating and binding the pore in its closed configuration (tonic block, TB) or by accessing and blocking the cavity once it has opened (use-dependent block, UDB)^32^. These agents are known to exert a more potent inhibitory effect on Na^+^ current during repetitive depolarization which accumulates the number of pores in their open or inactivated state (UDB), than during single stimuli from resting conditions which leave many channels still closed (TB)^4^. In each instance, the potency of drug block can be quantified by the half maximal inhibitory concentration (IC_50_), measuring the concentration of drug necessary to achieve 50% inhibition of sodium current^33^. To validate the tonic and use-dependent block properties of mexiletine using our automated planar patch clamp system, the mexiletine response of WT-only channels was measured at 0 Hz, 10 Hz and 20 Hz. As expected, the IC_50_ curves become increasingly left shifted as the frequency of stimulation is increased from 0 Hz (TB) to 10 and 20 Hz (UDB) *(Figure 1D).* The dose response for mexiletine was then recorded in R34C, H558R, S524Y and S1103Y mutant Nav1.5 channels and compared to wild type Nav1.5 at 0 Hz (*Figure 1E*) and 10 Hz *(Figure 1F).* The mexiletine dose-response curves for S1103Y (IC_50_ = 57.2 ± 14.3 μM; 13.4 ± 2.7 μM at 0 Hz and 10 Hz, respectively) show a substantial left shift compared to WT (IC_50_ = 148.6 ± 27.8 μM; 31.7 ± 4.4 μM). The measured differences in IC_50_ between WT and S1103Y represent a ∼2.6-fold more potent effect of mexiletine resting-state block of S1103Y (p = 0.006) and a ∼2.4-fold increase of drug effect at 10 Hz (p = 0.014). Interestingly, the other variants show similar or slightly right-shifted concentration curves compared to WT, indicating no significant effect on tonic or use-dependent inhibition of these channels. The mexiletine IC_50_ was calculated for wild type and variant Nav1.5 channels at each frequency tested with 95% confidence intervals (*Table 1*). To better understand whether the enhanced drug block in the S1103Y polymorphism is maintained with other class 1b antiarrhythmics, we also measured the lidocaine dose response across the four variants. To our surprise, S1103Y cells did not exhibit significant changes in IC_50_ for either tonic or use-dependent stimulation, while the common H558R variant showed a left-shifted pharmacology curve compared to WT (*SI Figure #1*). While more investigation is needed to elucidate the mechanism, our data is consistent with the observed clinical inconsistency in drug effectiveness among arrhythmic patients. These results suggest that lidocaine and mexiletine, although analogous, interact differently with the cardiac sodium channel.

**Figure 1:**
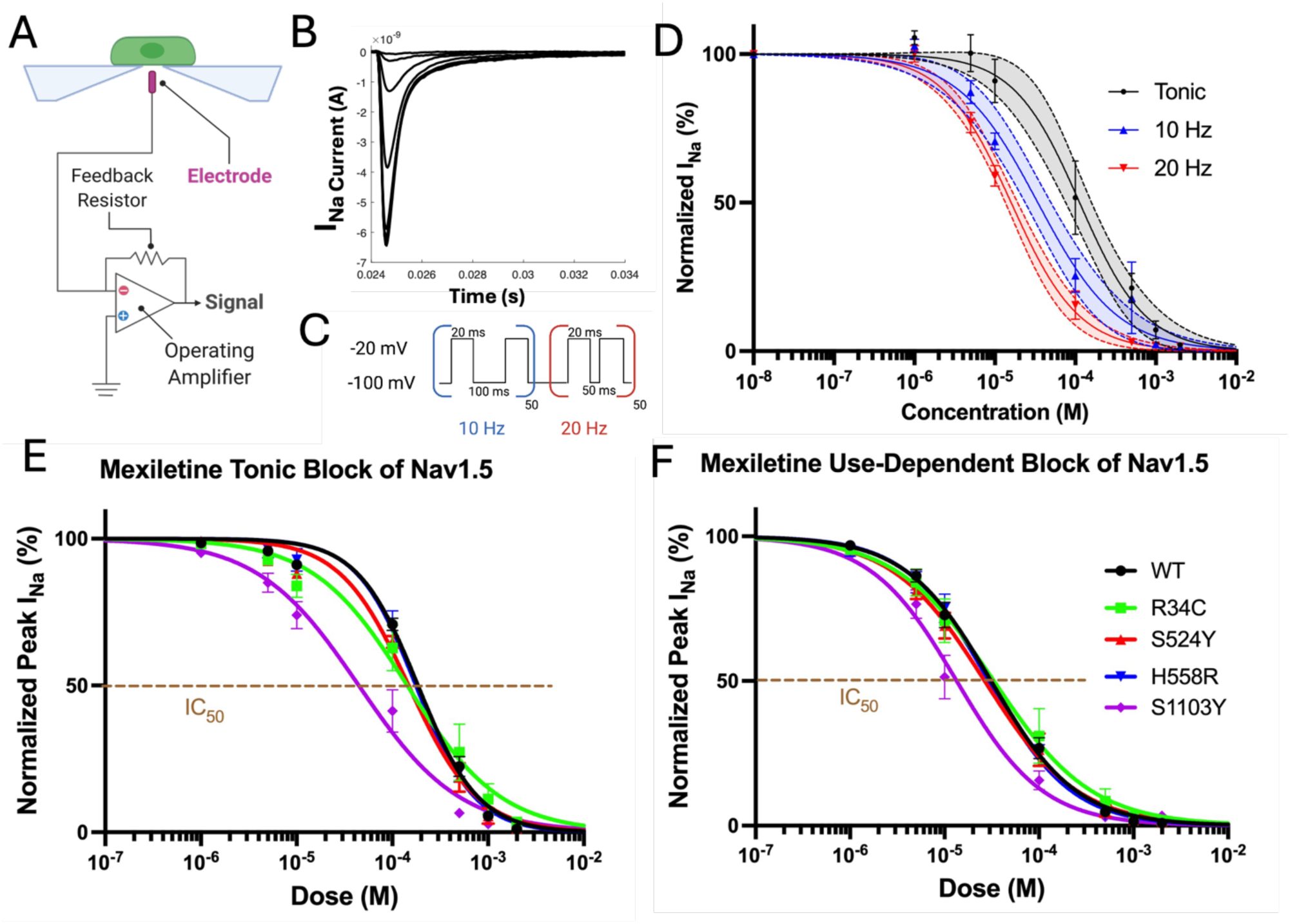
Automated planar patch clamp recordings of the mexiletine block of Nav1.5 currents in transiently transfected HEK293 cells. A) Patchliner schematic. B) Representative I_Na_ trace. C) The protocol used to measure UDB consists of a train of 50 test pulses to −20mV at varying frequencies. D) As the frequency of stimulation is increased, the mexiletine dose response becomes increasingly left-shifted. E) Mexiletine tonic block measured at 0Hz frequency. The S1103Y IC_50_ (44.6±10.2 µM) is significantly left-shifted compared to WT (IC_50_: 185.1±22.4 µM), p=0.006. The other variants do not exhibit a significant effect. B) Mexiletine use-dependent block measured at 10Hz frequency. The S1103Y IC_50_ (13.4±2.7 µM) is significantly left-shifted compared to WT (IC_50_: 31.1±4.4 µM), p=0.014. The other variants do not exhibit a significant effect. Sample size is 8-12 cells per variant.

**Table 1:** Mexiletine IC_50_ values for WT and variant Nav1.5 at 0 Hz and 10 Hz (95% CI).

|  | IC<sub>50</sub> at 0 Hz | IC<sub>50</sub> at 10 Hz |
| --- | --- | --- |
| **WT** | 148.6 ± 27.8 μM | 31.7 ± 4.4 μM |
| **R34C** | 143.1 ± 44.1 μM | 33.3 ± 9.4 μM |
| **S524Y** | 147.9 ± 24.6 μM | 26.6 ± 4.7 μM |
| **H558R** | 176.5 ± 26.7 μM | 29.4 ± 4.7 μM |
| **S1103Y** | 57.2 ± 14.3 μM | 13.4 ± 2.7 μM |

While results in our heterologous HEK cells system suggest an increased interaction with mexiletine in the S1103Y variant, we pursued further analysis in a physiologically relevant model more consistent with native myocyte physiology. To that end, CRISPR/Cas9 technology was used to engineer a line of hiPSCs expressing the S>Y mutation at the 1103 locus, using a well-characterized wild type hiPSC line (derived from a healthy male donor, WTc-11) as background. Cardiomyocytes were subsequently developed through established in-vitro differentiation protocols.

Simultaneously, a black female patient who was heterozygous for S1103Y (S1103Y^+/-^) was enrolled in the Mexiletine for Ventricular Tachyarrhythmia (MVET) clinical study at Washington University School of Medicine. This individual, referred to as MVET-55, presented to the clinic with end stage non-ischemic cardiomyopathy post-left ventricular assist device placement and a history of severe ventricular arrhythmias post implantable cardiac defibrillator who had breakthrough arrhythmia while prescribed chronic amiodarone therapy. The patient was prescribed mexiletine as adjunctive antiarrhythmic therapy, but due to progressive heart failure, underwent orthotopic heart transplant. PBMC-derived iPSCs were obtained from this patient to validate the WTc-derived S1103Y iPSC model. The patient heterozygous line was then used as background to engineer corresponding S1103Y^-/-^ and S1103Y^+/+^ myocytes. Taken together, these five lines of iPSC-derived cardiomyocytes allow the investigation of the specific role of the S>Y polymorphism in altering response to mexiletine, while keeping patient background (either WTc or MVET-55) as a fixed variable in each group.

Since MVET-55 cells were developed for the first time in this study, confocal microscopy was used to characterize the baseline morphological and structural properties. Consistent with previous findings in native and iPSC-derived cardiomyocytes, cells showed well-defined striations of the z-discs (α-actinin; blue) and established cell-to-cell connections (N-cadherin in adherens junctions; yellow)^34,35^. Nav1.5 (purple) was shown to colocalize with the z disc (*Figure 2A-C*) and intercalated disc (*Figure 2D-F*) nanodomains in all three hiPSC-derived cardiomyocyte lines.

**Figure 2:**
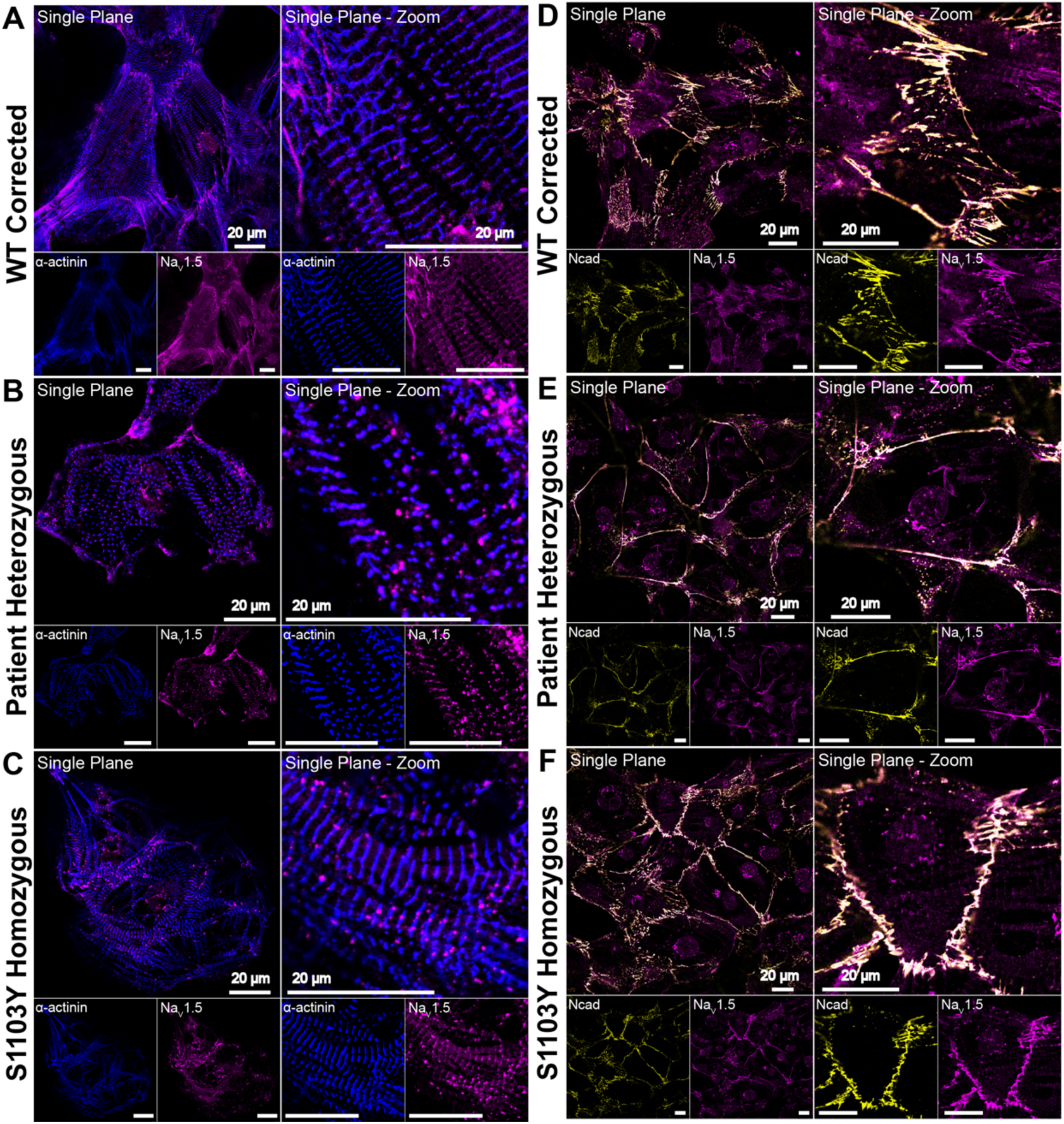
Na_V_1.5 localization within iPSC-derived cardiomyocyte nanodomains. Na_V_1.5 (purple) colocalizes with α-actinin at the z-discs **(A-C,** blue) and with N-cadherin at the intercalated discs **(D-F,** yellow) in WT corrected S1103Y^-/-^ **(A, D)**, patient heterozygous S1103Y^+/-^ **(B, E)**, and homozygous S1103Y^+/+^ **(C, F)** cardiomyocytes. Single plane images were presented as whole cells and zoomed in views.

The mexiletine dose response was characterized in each line under both tonic and use-dependent stimulation. In the WTc-background lines (*Figure 3*, top panel), the S1103Y^+/+^ dose response exhibited a significant left shift compared to WT^+/+^ in both tonic block (S1103Y IC_50_: 7.08±1.8 µM; WTc IC_50_: 100.3±22.9 µM; p=0.0005) and use-dependent block at 10^+/+^ Hz frequency (S1103Y IC_50_ (4.2±1.0 µM; WTc IC_50_: 24.4±2.7µM; p=0.0004), consistent with our previous HEK cell data (*Figure 3B,C*). Interestingly, the same pharmacology study in MVET-55 background lines demonstrated that S>Y mutation of both alleles at locus 1103 is required to observe a significant left shift in the mexiletine dose response, for either tonic or use-dependent block. The dose response curves revealed an IC_50_ of 37.8±7.8 µM for the S1103Y^-/-^ line at 0 Hz, compared to 12.2±3.3 µM for the S1103Y^+/+^ line, indicating a significantly increased potency of mexiletine (p=0.018). Similar, the left shift was maintained at 10 Hz, with an IC_50_ of 23.9±5.0µM in S1103Y^-/-^ and 8.2±2.1 µM in S1103Y^+/+^ (p=0.042). The patient-derived heterozygous variant S1103Y exhibited a dose response similar to the WT corrected line, with IC_50_ of 39.7±8.1 µM at 0Hz and 12.7±2.1 µM at 10Hz (p=0.98 and p=0.66, respectively), (*Figure 3E,F*). Taken together, the pharmacology results discussed above imply an enhancement of the mexiletine block of peak Nav1.5 current in S1103Y homozygous cardiomyocytes.

**Figure 3:**
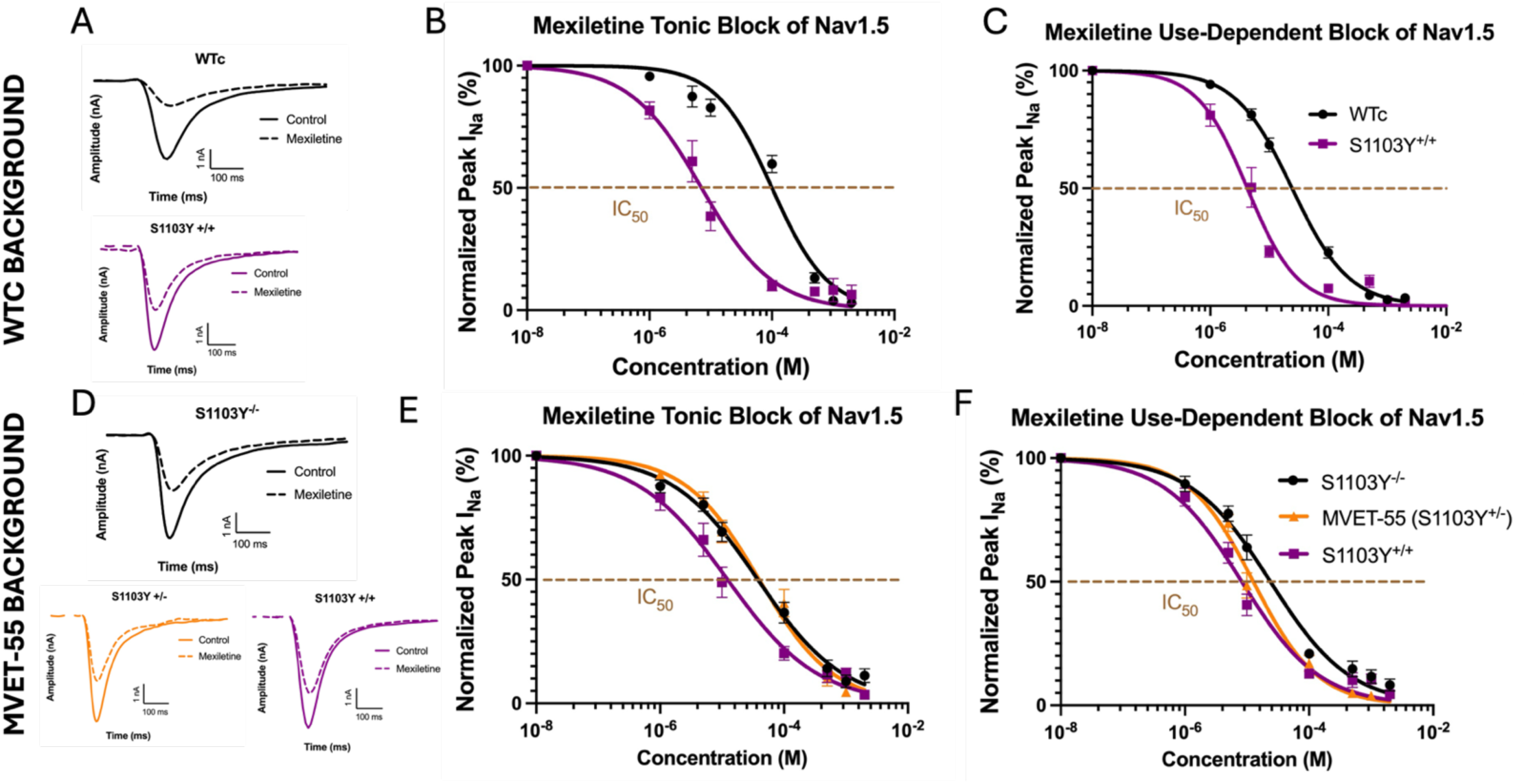
Mexiletine block of Nav1.5 currents in iPSC-CMs lines with WTc or MVET-55 background. *Top panel:* the S1103Y homozygous line was CRISPR-engineered from the WTc line. A) Representative I_Na_^+/+^ traces before and after mexiletine. B) Mexiletine tonic block measured at 0Hz stimulation. The S1103Y^+/+^ IC_50_ (7.08±1.8 µM) is significantly left-shifted compared to WTc (IC_50_: 100.3±22.9 µM), p=0.0005. C) Mexiletine use-dependent block measured at 10Hz frequency. The S1103Y^+/+^ IC_50_ (4.2±1.0 µM) is significantly left-shifted compared to WT corrected (IC_50_: 24.4±2.7µM), p=0.0004. *Bottom panel:* the WT corrected and S1103Y homozygous lines were CRISPR-engineered from the patient-derived MVET-55 line (S1103Y heterozygous). D) Representative I_Na_ traces before and after mexiletine. E) Mexiletine tonic block measured at 0Hz frequency. The S1103Y^+/+^ IC_50_ (12.2±3.3 µM) is significantly left-shifted compared to WT corrected (IC_50_: 37.8±7.8 µM), p=0.018. S1103Y^+/+^ (IC_50_: 39.7±8.1 µM) exhibits a dose response similar to WT corrected (p=0.98), while it is significantly right-shifted compared to S1103Y (p=0.018). F) Mexiletine use-dependent block measured at 10Hz frequency. The S1103Y^+/+^ IC_50_ (8.2±2.1 µM) is significantly left-shifted compared to WT corrected (IC_50_: 23.9±5.0µM), p=0.042. The S1103Y IC_50_ (13.7±2.2 µM) does not exhibit a significant effect compared to WT corrected (p=0.66) or S1103Y^+/+^ (p=0.13). Sample size is 8-13 cells per line. Nav1.5 currents are recorded with automated planar patch clamp.

To further investigate the drug response, we measured the action potential duration at 90% repolarization (APD_90_) across the MVET-55 lines, comparing control and mexiletine-treated cells. Previous studies indicate that mexiletine exerts QTc shortening antiarrhythmic properties by preferentially inhibiting the late component of the sodium current (rather than peak), which is often associated with proarrhythmic APD_90_ prolongation in LQTS disease states^36,37^. Various reports also suggest that in wild type, healthy cells, mexiletine does not substantially alter the repolarization phase of the action potential.^38–40^ Consistent with this finding, optical electrophysiology demonstrated that the APD_90_ was not significantly altered by acute mexiletine application in S1103Y^-/-^ and S1103Y^+/-^ cells, which also did not exhibit an altered pharmacological response to the drug in our dose-response experiments. Interestingly, the S1103Y^+/+^ cells responded differently to mexiletine, exhibiting pronounced APD_90_ prolongation, a phenotype that is connected to arrhythmia propensity and not the expected outcome of a Class 1b antiarrhythmic drug (*Figure 4B)*. This unanticipated APD_90_ prolongation after acute mexiletine treatment was robustly recapitulated in multiple S1103Y^+/+^ lines, including one clone of the WTc-11 derived homozygous line and two different clones of the MVET-55 derived homozygous line (4B3 and 3E7) (*SI Figures #2 and #3*).

**Figure 4:**
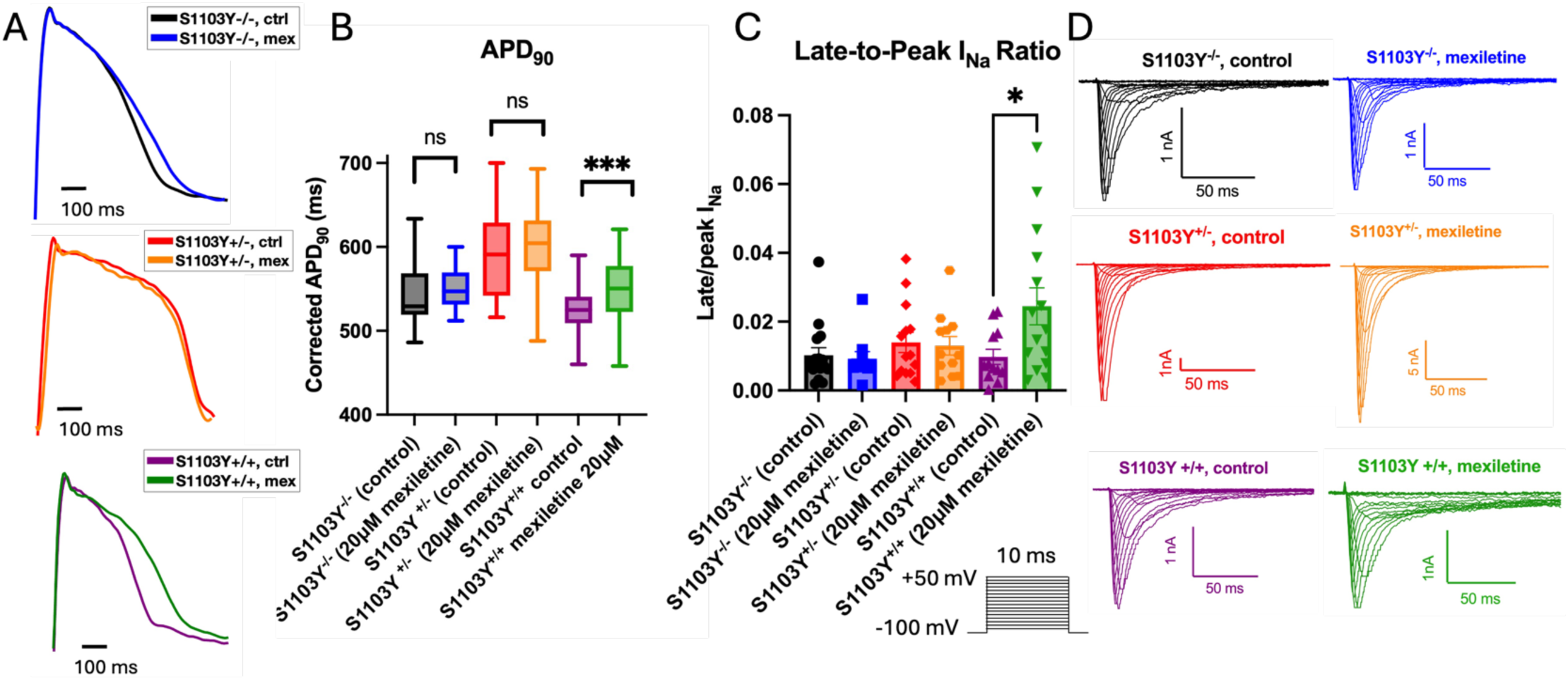
Action potential duration (APD) and late sodium current measurements in MVET-55 background iPSC-CM lines. A) Representative APD_90_ traces measured at 1Hz in control and mexiletine-treated cells. 20μM mexiletine was applied 15 minutes before recording. The Savitzky-Golay filter was applied to all representative traces. B) Mexiletine significantly prolongs the APD_90_ (+175.8ms) in the S1103Y homozygous line (p < 0.001) compared to control. The WT corrected and S1103Y heterozygous lines do not exhibit significant changes in APD_90_ after mexiletine treatment. C) Late-to-peak sodium current ratio in control and mexiletine-treated cells measured at −20mV. The S1103Y homozygous line shows a significant increase in late-to-peak I_Na_ ratio (+0.0147, p = 0.02) after mexiletine treatment. No changes are observed in the WT corrected and S1103Y heterozygous line. D) Representative sodium current traces in control and mexiletine treated cells in WT corrected (top row), S1103Y heterozygous (patient line, middle row), or S1103Y homozygous iPSC-CMs, with recording voltage protocol in inset.

To elucidate the mechanism of APD_90_ prolongation, automated patch clamp was used to record Nav1.5 currents and isolate the persistent component, as measured at 20ms from the peak current measurement and reported as the ratio between late and peak values. Consistent with the APD_90_ prolongation observed in mexiletine-treated S1103Y^+/+^ cells, Nav1.5 recordings revealed an increase in late sodium current compared to control, while the late current in S1103Y^-/-^ or S1103Y^+/-^myocytes did not vary after drug treatment (*Figure 4C*). Taken together, the enhanced dose response for peak current block and the paradoxical increased late current in mexiletine-treated S1103Y^+/+^ cells indicate that this variant, when present in both alleles, exhibits a peculiar response to the drug, preferentially inhibiting the peak sodium current rather than the late, pathogenic component.

To further understand if the Nav1.5 kinetics are altered differently by mexiletine across the MVET-55 lines, we used automated patch clamp to characterize the activation, inactivation and recovery properties (*Figure 5*). While mexiletine did not alter peak sodium current in S1103Y^-/-^ cells, it slightly reduced current density in the S1103Y heterozygous and homozygous myocytes (*Figure 5A*), although the change was not significant (p=0.55 and 0.72, respectively). No substantial differences were observed in the conductance-voltage or inactivation traces after drug treatment (*Figure 5B*). Mexiletine has been previously shown to slow down both phases of channel recovery from inactivation, which was also recapitulated across the three MVET-55 lines (*Figure 5C*). While the fast component was substantially slower, the slow phase reduction was not significant. Overall, the Nav1.5 gating kinetics were not dramatically altered in the S>Y variant lines, in the presence or absence of mexiletine (*Table 2*). This finding is consistent with the non-pathogenic nature of the S>Y mutation, which is classified as a common polymorphism present in a large portion of African Americans (∼13%), and thus not expected to directly cause arrhythmogenic alterations to sodium channel kinetics.

**Figure 5:**
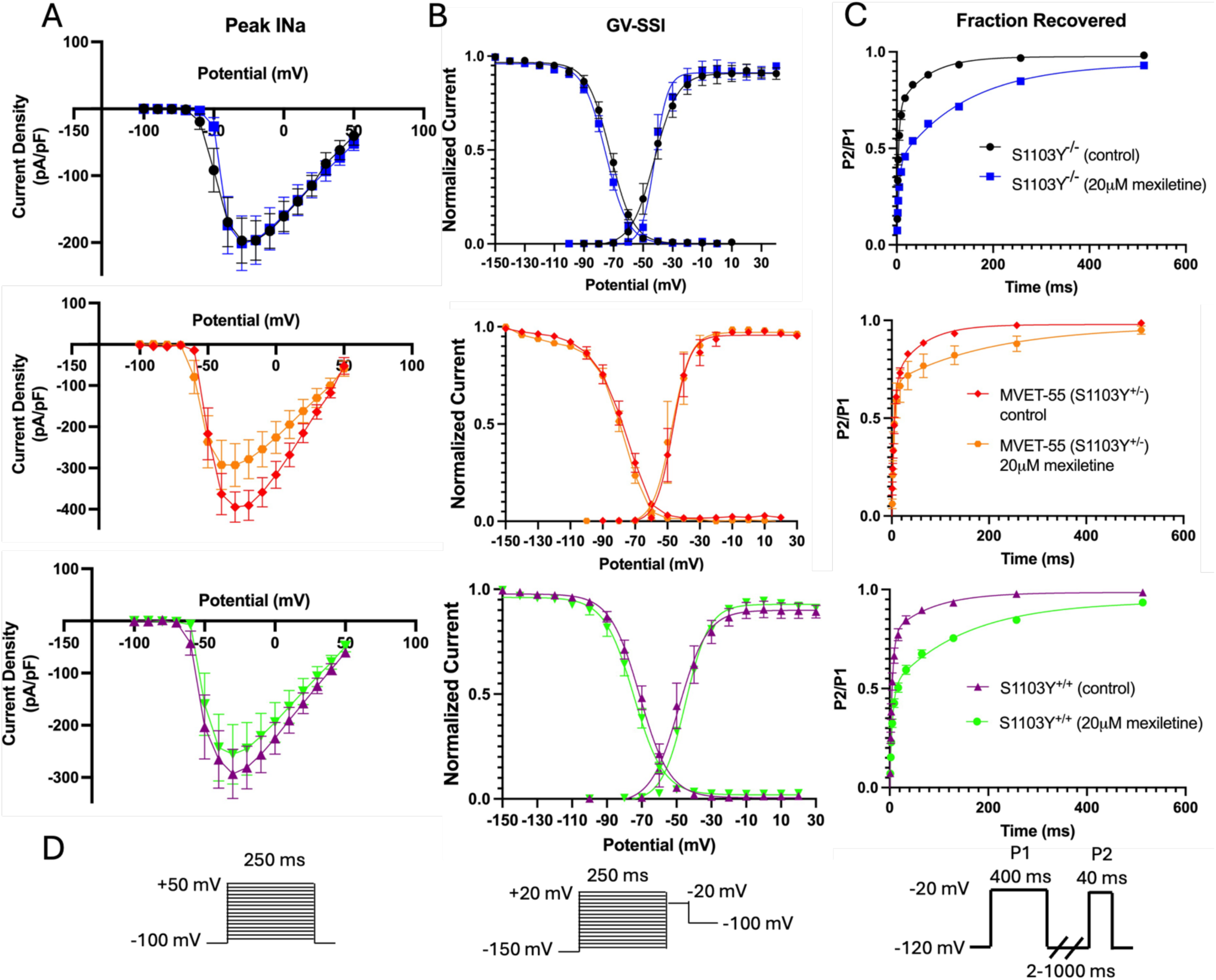
Nav1.5 kinetics in control and mexiletine-treated iPSC-CMs. A) Acute mexiletine (20µM) application does not significantly reduces peak I^Na^ in the S1103Y^-/-^, S1103Y^+/-^ or S1103Y^+/+^ lines (p = 0.99, 0.55, 0.72 respectively). B) The conductance-voltage and steady-state inactivation curves are not significantly different across the S1103Y^-/-^, S1103Y^+/-^, and S1103Y^+/+^ lines in control vs mexiletine-treated cells (GV V1/2: p > 0.99. SSI V^1/2^: p > 0.99). C) Mexiletine slows down the fast phase of recovery in S1103Y^-/-^, S1103Y^+/+^, and S1103Y cells. τ_10_ p-values = 0.0001, 0.04, 0.02, respectively. The slow phase of recovery, although slightly slower with mexiletine, is not significantly different. τ_1_ p-values > 0.99. D) Voltage protocols for Nav1.5 activation, inactivation and recovery.

**Table 2:**
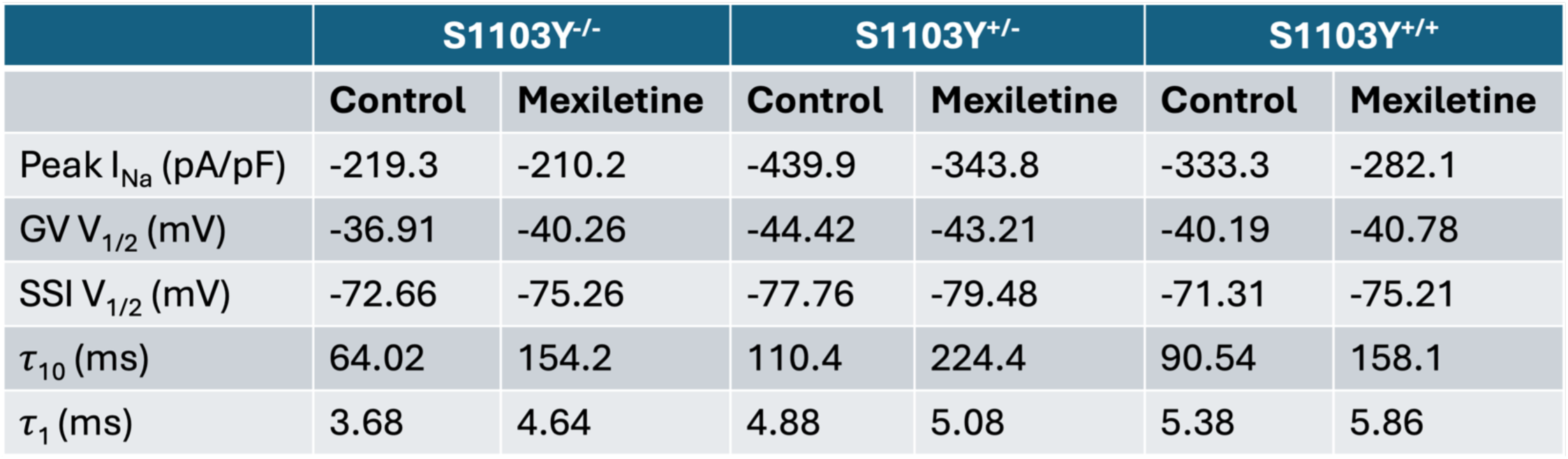
Summary of Nav1.5 kinetics for peak I_Na_, half-maximal voltage of activation and inactivation, and fast and slow recovery time constants.

## Discussion

The present study probes the link between genetic variability in the cardiac sodium channel and antiarrhythmic therapy outcomes. We demonstrate that common SCN5A polymorphisms, while not evoking major electrophysiological changes to background channel function, may unmask a proarrhythmic phenotype and alter the pharmacological response to class 1b antiarrhythmics. Although previous studies have reported that the S1103Y variant, commonly found in African-descendent populations, may be associated with increased arrhythmogenic risk in individuals already suffering from heart disease^41–43^, our results reveal that this single point polymorphism elicits an unexpected response to mexiletine. We also show that S>Y mutation of one allele is not sufficient to alter the drug phenotype. Rather, only S1103Y homozygous myocytes exhibit previously unreported, likely arrhythmogenic changes to late sodium current and APD_90_ after mexiletine treatment.

### Physiological relevance

Many mutations have been identified in the SCN5A gene. Among these, some are pathogenic and have a direct association with cardiac arrhythmias, including long QT syndrome type 3 (LQTS3), Brugada syndrome, and cardiac conduction diseases^44^. Others are classified as benign and are present with relatively high frequency in the general population without an obvious correlation with cardiac pathology^45,46^. Among these, H558R is the most common sodium channel polymorphism, widely detected in all ethnic groups (29% of Blacks, 20% of Whites, 23% of Hispanics, and 9% of Asians)^21^. Other polymorphisms only appear in specific ethnicities. S524Y and S1103Y are detected exclusively in populations of African descent and have a prevalence of 6% and 13%, respectively. R34C is found predominantly among Blacks (9%), although it is also present among Asians (3%)^21^. These common variants are positioned in the intracellular portion of the channel, with R34C being located in the N-terminus; S524Y and H558R in the DI-DII linker; and S1103Y in the DII-DIII linker^21^. Notably, the DI-DII and DII-DIII cytoplasmic regions are still absent from solved Nav1.5 structures and their roles remain elusive, indicating that the functional significance of variants in these regions is largely unknown^47^. Previous reports suggest that carrying the S1103Y allele may increase arrhythmia propensity in black individuals with heart failure, although the variant itself is not generally considered pathogenic^42^. Thus, identifying unique genotypes that may render popular antiarrhythmics *proarrhythmic* in susceptible genetic backgrounds for common arrhythmia syndromes (e.g. advanced heart failure and ischemic cardiomyopathy) is critically important.

Cardiac arrhythmias are observed with high prevalence in patients across all genetic backgrounds. However, most class 1b antiarrhythmic drugs, including lidocaine and mexiletine, were developed in the 1980s at a time when genetic variation was not recognized as an important selection criterion during clinical trials^48,49^. To date, antiarrhythmic therapy for common, acquired heart syndromes does not consider the variability in nucleotide sequences encoding the cardiac ion channels that can trigger those arrhythmias. As a result, drug therapy remains inconsistent among patients and often cause adverse effects that increase toxicity and may result in increased proarrhythmic risk^50^. Lidocaine and mexiletine work by blocking sodium channels, binding primarily to the open-inactivated state and reducing cellular excitability, which can disrupt the propagation of re-entrant rhythms^51,52^. Lidocaine is very effective in terminating ventricular tachycardia after acute myocardial infarction, but it requires an intravenous formulation and is thus only used in the acute, inpatient setting^16,17^. Mexiletine can be administered orally, but its efficacy is demonstrated only for a small subset of patients. Previous cardiac catheterization studies reported that mexiletine successfully prevented arrhythmia induction in the context of programmed electrical stimulation in ∼20% of patients^53–55^. Thus, there is a clinical need to develop new, individualized therapeutic approaches that are highly effective for patients with different genetic backgrounds and are suitable for at-home and long-term treatment.

### Comparing the mexiletine response in common SCN5A polymorphisms

Our work establishes that a single point polymorphism in SCN5A, S1103Y, detected exclusively among populations of African ancestry, paradoxically alters mexiletine response compared to wild type channels. The presence of the S1103Y allele, whether in a heterologous mammalian system or in hiPSC-cardiomyocytes, robustly left shifted the dose response to mexiletine. The substantial effect was observed for both tonic and use-dependent inhibition, suggesting that S1103Y patients may experience increased peak sodium current block with mexiletine therapy both during unstimulated periods of rest (tonic block) as well as during episodes of heightened cardiac activity (use-dependent block), such as during ventricular tachycardia. R34C, S524Y and H558R did not significantly affect drug binding in HEK cells. Interestingly, the same S1103Y variant did not exhibit a left-shifted response when treated with lidocaine, which is structurally and functionally similar to mexiletine, with IC_50_ values in the same range as wild type (*SI Figure #1)*.

While the left shift in dose response implies an increased interaction with mexiletine in the S1103Y-carrying Nav1.5 gene, the effect only demonstrates enhanced drug block of the *peak* component of the sodium current (I_Na,P_). Nav1.5-associated arrhythmic disorders like LQTS3, however, are connected to an increase in the late component of sodium current (I_Na,L_), which prolongs the action potential duration and is seen on electrocardiogram as a prolonged QT interval. Thus, the therapeutic potential of class 1b antiarrhythmics lies in their ability to preferentially block I_Na,L_ over I_Na,P_. Interestingly, our automated patch clamp recordings in iPSC-derived cardiomyocytes revealed an unexpected finding, with S1103Y cells presenting increased I_Na,L,_ when treated with mexiletine. While I_Na,L_ is normally 0.1 to 1% of I_Na,P_ in wild type myocytes, the ratio was ∼2.5% in cells carrying the mutation when mexiletine was added. This proarrhythmic effect, however, was only observed in S1103Y homozygous cells, indicating that presence of both S>Y alleles is necessary to perturb the mexiletine response. Previous reports have demonstrated an α-subunit multimerization mechanism, where individual subunits physically interact with one another and function as a coupled dimer via protein 14-3-3 mediation^56^ to significantly alter channel function. Although further investigation is needed, this paradigm of sodium channel assembly may explain why only the homozygous variant exhibits an altered response to mexiletine, while the heterozygous counterpart maintains a drug phenotype similar to wild type channels.

To further investigate the translational significance of the S1103Y polymorphism, we measured action potential duration at 90% repolarization in the presence or absence of mexiletine. A prolongation of APD_90_ is considered proarrhythmic, as it provides an opportunity for calcium channels to prematurely recover from inactivation and trigger early after-depolarization events, which can lead to Torsade de pointes^57^. An increased I_Na,L_ is a substrate for APD_90_ prolongation and thus arrhythmogenic. Consistent with our late current measurements, optical microscopy experiments revealed that S1103Y^+/+^ myocytes exhibit a prolonged APD_90_ when treated acutely with mexiletine. Taken together, our results suggest that the mexiletine response in S1103Y patients may be opposite to what physicians would expect, reversing the canonical drug phenotype and leading to enhanced block of I_Na,P_ while also triggering pathogenic I_Na,L._

### S1103Y does not substantially alter Nav1.5 kinetics

Genetic variation is often the result of random mutations that occur during DNA replication and are subsequently passed on to the offspring. Evolutionarily, common sodium channel polymorphisms had an opportunity to become prevalent in the general population thanks to their benign phenotype. Cardiac sodium channels are essential in maintaining myocyte excitability and normal sinus rhythm by underlying the initiation (phase 0) and rapid firing of the action potential. Therefore, it is expected that common Nav1.5 variant proteins evolved to preserve crucial Nav1.5 kinetics, maintaining their wild type-like function. To test this hypothesis, automated patch clamp was used to characterize the activation, inactivation and recovery parameters of Nav1.5 channels in cells with identical background, derived from the MVET-55 patient, except at the 1103 locus. The iPSC-derived myocytes either carried the original S1103Y^+/-^ variant or were CRISPR-engineered to express the S1103Y^+/+^ double allele or corrected to the wild type-like phenotype S1103Y^-/-^. As anticipated, no significant changes were recorded across the three cell lines, even in the presence of mexiletine, when comparing the V_1/2_ of activation, V_1/2_ of inactivation, and recovery time constants. The pronounced differences in dose response and the simultaneous conservation of Nav1.5 kinetics suggest that these channels evolved to retain their fundamental cellular function but likely did not evolve to preserve the drug binding characteristics. Since class 1b antiarrhythmics preferentially bind to the inactivated state, it is possible that the S1103Y variant alters the conformation, rather than the kinetics of rearrangement, of the drug-binding pocket. While further structural studies and molecular docking simulations would be necessary to elucidate the exact binding properties, our data highlights the importance of understanding how genetic variation in key cardiac genes affects antiarrhythmic drug response. In light of our findings, we suggest caution when prescribing mexiletine to S1103Y patients. The drug response is likely to be opposite of the anticipated outcome, evoking increased late sodium current and further prolonging the APD_90_, which may carry serious, potentially life-threatening consequences in individuals already at risk for cardiac arrhythmias.

### Mexiletine access to its receptors occurs via a dual mechanism

The canonical antiarrhythmic drug receptor in sodium channels is located intracellularly in the central cavity of the pore, nested between residues from the S6 segments of all four domains^58^. Previous studies have described two distinct pathways through which antiarrhythmics can interact with the receptor. One is a hydrophilic pathway which depends on the state of the channel, as drugs can dock to the receptor from the intracellular side after permeating the open pore, in what is known as open-state block^32,59^. The other is a hydrophobic pathway, which allows lipid-soluble compounds to diffuse through the membrane and bind directly to the receptor. This latter mechanism is thought to be mediated by fenestrations, which are lateral openings in the sodium channel pore, first discovered in the ancestral bacterial sodium channel NavAB and conserved over billions of years of evolution from prokaryotes to eukaryotes^60,61^. Fenestrations have been suggested as a possible pathway for antiarrhythmic drugs because they provide hydrophobic tunnels that allow direct access from the lipid membrane to the receptor site in the pore^62^. In recent years, a second drug binding site has been suggested, located in the lateral portion of the pore in the DIII-DIV fenestration^20^. Most antiarrhythmic drugs exhibit a significant dependence on the pH of the local environment, as evidenced by the presence of an uncharged and charged moiety^63^. Mexiletine and lidocaine both have an ionizable amine group which, in its protonated state, confers a positive charge to the molecule. Lidocaine has a pKa of 7.8^64^, indicating that at physiological pH it is in part uncharged and hydrophobic^11^. Mexiletine has a pKa of 9.52^65^, indicating that it is mostly ionized and hydrophilic at physiological pH^19^. It is well recognized that heart disease disrupts blood pH, which in turn affects cardiac ion channel function and can increase proarrhythmic susceptibility^66^. Previous literature has suggested that lidocaine is more lipid-soluble and can interact with the Nav1.5 pore through fenestrations^60,67^. Conversely, the more hydrophilic mexiletine displays a drug-receptor interaction that depends on the state of the channel. Thus, variants that alter sodium current kinetics would also alter this receptor-drug interaction, explaining why only a subset of individuals are responsive to mexiletine, while the lidocaine effect is more consistent across patients. Interestingly, we have identified a variant, S1103Y, which exhibits a pronounced and unexpected response to mexiletine, despite a lack of effect on Nav1.5 kinetics. While activation, inactivation and recovery are unchanged, the late sodium current is significantly increased in S1103Y myocytes. The 1103 locus in the DII-DIII linker does not suggest a direct interaction with either binding site in the pore or in the DIII-DIV fenestration. Taken together, these observations suggest an allosteric mechanism that changes the conformation of the Nav1.5 pore, generating a leaky channel when mexiletine is present. Future structural studies with Nav1.5 in the inactivated state and mexiletine bound to the receptor or fenestration may help elucidate the mechanism underlying this peculiar drug response in S1103Y channels.

### Patient Genotyping and iPSC Modeling to Predict Pharmacological Responses

Our study introduces a novel, multidimensional approach to connect unique human genetic characteristics, iPSC-based in-vitro modeling, and high-throughput functional readouts of pharmacological responses on both molecular (ion channel) and cellular (action potential) human electrophysiology. Patient genotyping is already implemented in interventional cardiology^11^, to diagnose a wide array of inherited diseases^68^, and to guide targeted therapy in cancer and infectious diseases^69,70^. Combined with genomic engineering, we demonstrate here how iPSC-cardiomyocytes derived from individuals with genetic variants can be utilized to develop a physiologically relevant, patient-specific model that can subsequently be perturbed to predict a wide array of pharmacological responses in vitro, and thus improve efficacy of antiarrhythmic therapies. Although this study focuses on a cardiac sodium channel gene variant, S1103Y, and its ability to alter the mexiletine response in unsuspected ways, this framework can be applied to a wide variety of ion-channel associated mutations - within or outside cardiac genes - and to interrogate their specific effects on various pharmacological agents. Taken together, our precision medicine approach demonstrates its predictive value in disease mechanisms and drug responses, emphasizing the importance of integrating individual variability in clinical practice to achieve high therapeutic efficacy through personalized health applications.

## Limitations

CRISPR editing was used to genetically engineer the WTc and MVET-55 hiPSC lines to express the S1103Y point mutation in the Nav1.5 gene. While the same primers were used for both lines and standard quality control measures (i.e. sequencing, genotyping and karyotyping) were applied, we cannot exclude that the gene editing process affected Nav1.5 function, thus potentially confounding the drug response effects observed in our results. While we believe that hiPSC-derived cardiomyocytes represent a substantial improvement compared to heterologous mammalian systems to understand cardiac electrophysiology- HEK cells transfected with non-diseased Nav1.5 do not allow robust recordings of late sodium current- we also recognize that the current hiPSC-CM model is not a perfect proxy for ventricular myocytes due to immature electrical and structural composition^71,72^. Our studies demonstrate that genetic variability modulates antiarrhythmic drug responses. Nevertheless, further clinical studies are required to validate whether these perturbations are recapitulated at the patient level. To our knowledge, there are no reported electrocardiograms specific to the S1103Y homozygous patient population. Understanding whether these individuals experience unexpected pro-arrhythmic QTc prolongation after mexiletine treatment would corroborate our model and thus advocate for the use of genetic testing to inform arrythmia therapy.

## Conclusion

Our studies in hiPSC-CMs contribute to a deeper understanding of the role of genetic variability in key cardiac genes in modulating antiarrhythmic drug responses. We demonstrate here that a common point mutation in the Nav1.5 gene, S1103Y, robustly left shifts the mexiletine pharmacology curve compared to wild type under both tonic and use-dependent stimulation. While this effect implies a more pronounced block of the peak sodium current in this variant, we show that mexiletine unexpectedly increases the pathogenic late sodium current leading to proarrhythmic APD_90_ prolongation in S1103Y myocytes. Interestingly, we show that S1103Y homozygosity is required to induce these surprising alterations to the mexiletine response. Such information provides new insights into the inconsistencies in mexiletine effectiveness reported clinically and may be relevant for the improvement of patient-specific arrhythmia therapy.

## Data Availability Statement

The data that support the findings of this study are available from the corresponding author upon reasonable request and limitations with respect to patient privacy.

## Acknowledgments

We gratefully acknowledge the financial support provided by the National Heart Lung and Blood Institutes of Health (R01 HL148803 to Jonathan Silva) and the American Heart Association (24PRE1198562 to Martina Marras and 960621 to Jonathan Silva). Organizations that provided funding were not involved in study design, data collection, data analyses, data interpretation, manuscript preparation, or the decision to submit this article for consideration for publication.

## Supplementary Data

**SI Fig. #1:**
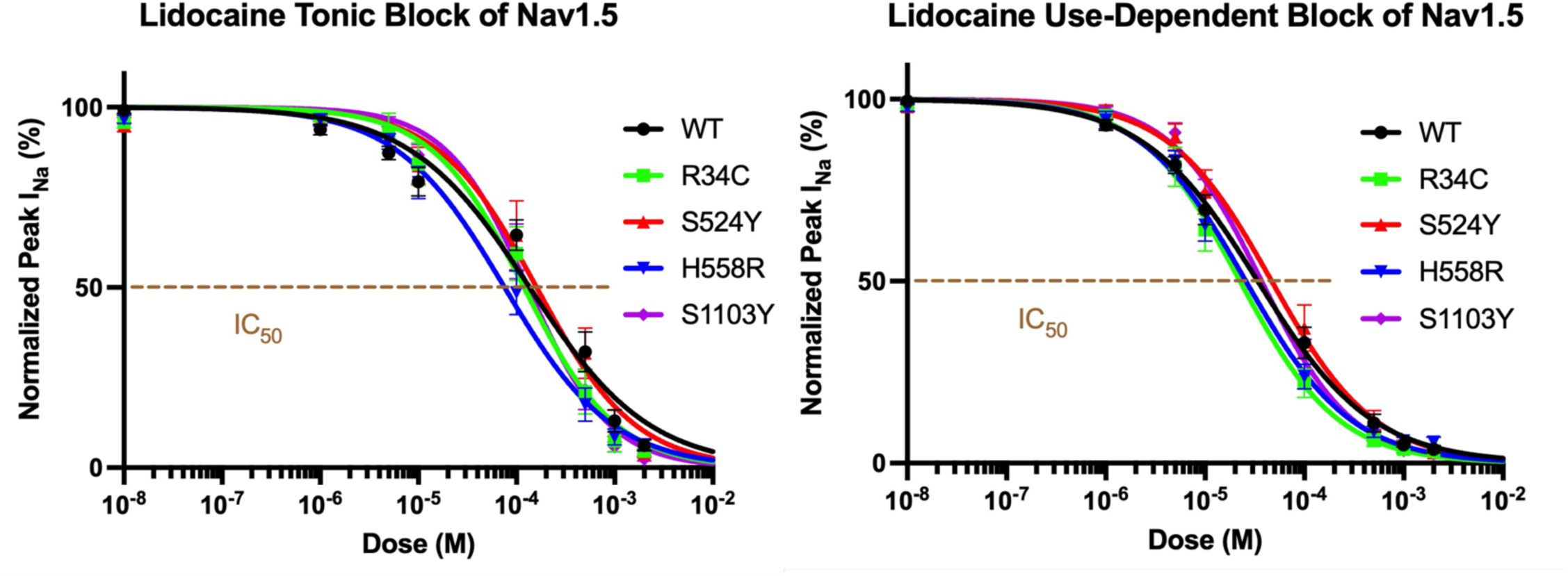
Lidocaine block of Nav1.5 currents in transiently transfected HEK293 cells. **A)** Lidocaine tonic block measured at 0Hz stimulation. The H558R IC_50_ (75.6±16.6 µM) is significantly left-shifted compared to WT (IC: 136.1±33.9 µM), p=0.003. The other variants do not exhibit a significant effect (p>0.05). B) Lidocaine use-dependent block measured at 10Hz frequency. The H558R IC_50_ (25.7±4.2 µM) is significantly left-shifted compared to WT (IC: 33.9±5.5 µM), p=0.03. The other variants do not exhibit a significant effect (p>0.05). Sample size is 8-12 cells per variant. Nav1.5 currents are recorded with automated planar patch clamp.

**SI Fig. #2.**
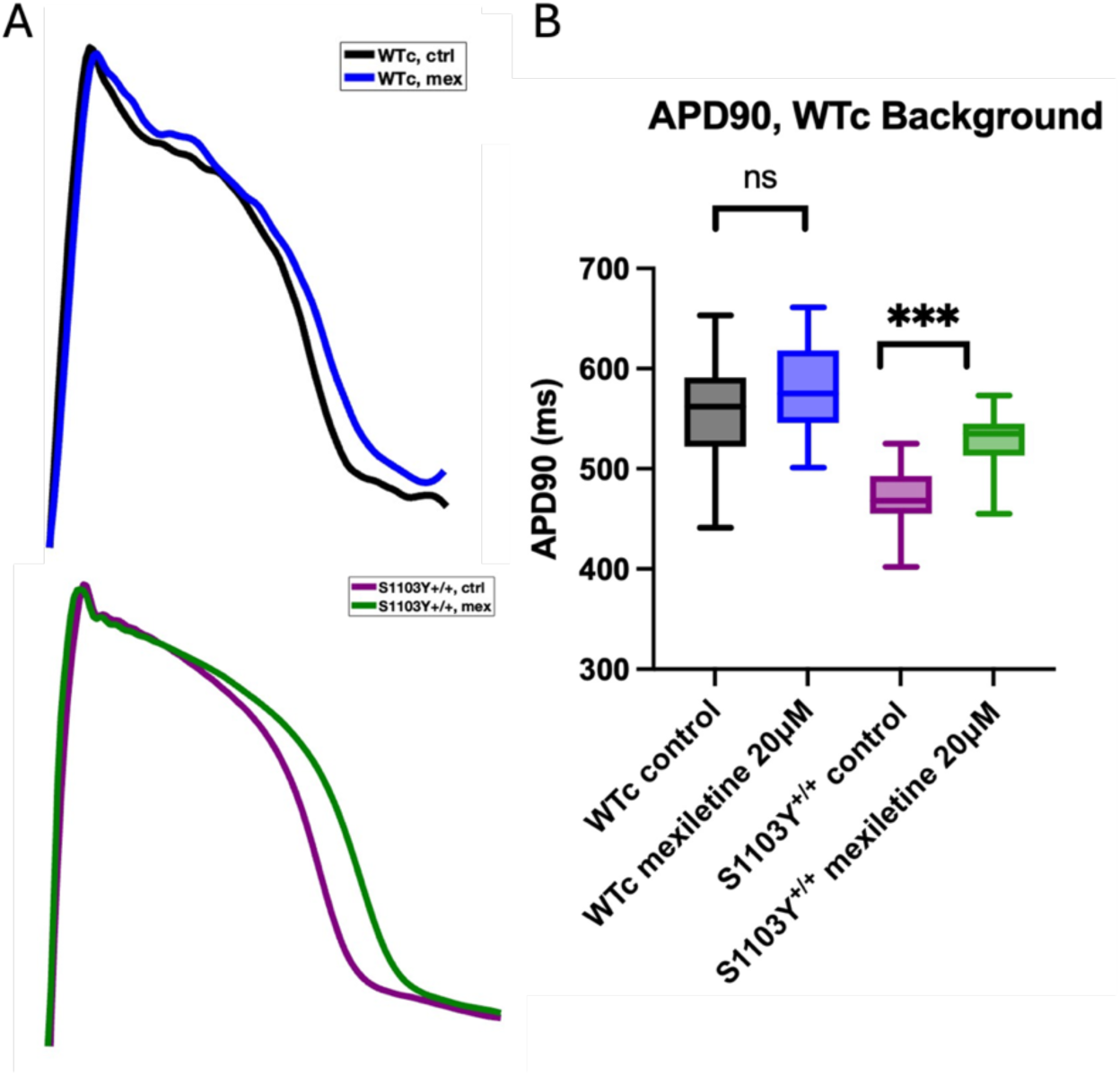
Action potential duration (APD) in WTc background iPSC-CM lines. A) Representative APD_90_ traces measured at 1Hz in control and mexiletine-treated cells. 20μM mexiletine was applied 15 minutes before recording. The Savitzky-Golay filter was applied to all representative traces. B) Mexiletine significantly prolongs the APD90 (+58.9ms) in the S1103Y homozygous line (p < 0.001) compared to control. The parent WTc line does not exhibit significant changes in APD_90_ after mexiletine treatment (p = 0.1).

**SI Fig. #3:**
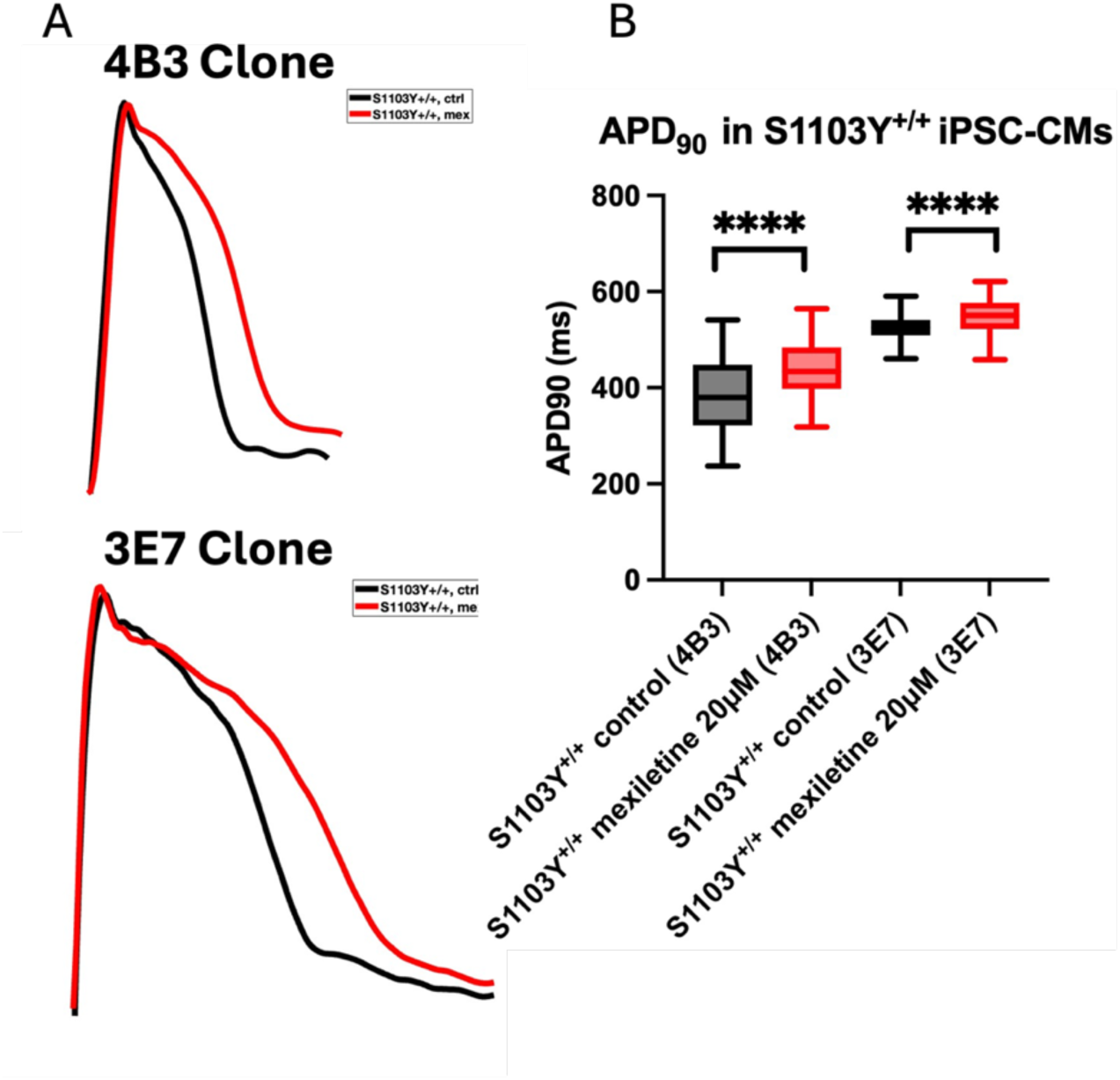
Action potential duration (APD) in two clones of S1103Y**^+/+^**, MVET-55 background iPSC-CMs. **A)** Representative APD_90_ traces measured at 1Hz in control and mexiletine-treated cells. 20μM mexiletine was applied 15 minutes before recording. The Savitzky-Golay filter was applied to all representative traces. B) Mexiletine significantly prolongs the APD_90_ by +48.6ms in the 4B3 clone (p < 0.001) and by +47.7ms (p = 0.008) in the 3E7 clone compared to control.

